## Supplementary Figures for "RNA polymerase II pausing is essential during spermatogenesis for appropriate gene expression and completion of meiosis"

**Figure S1**

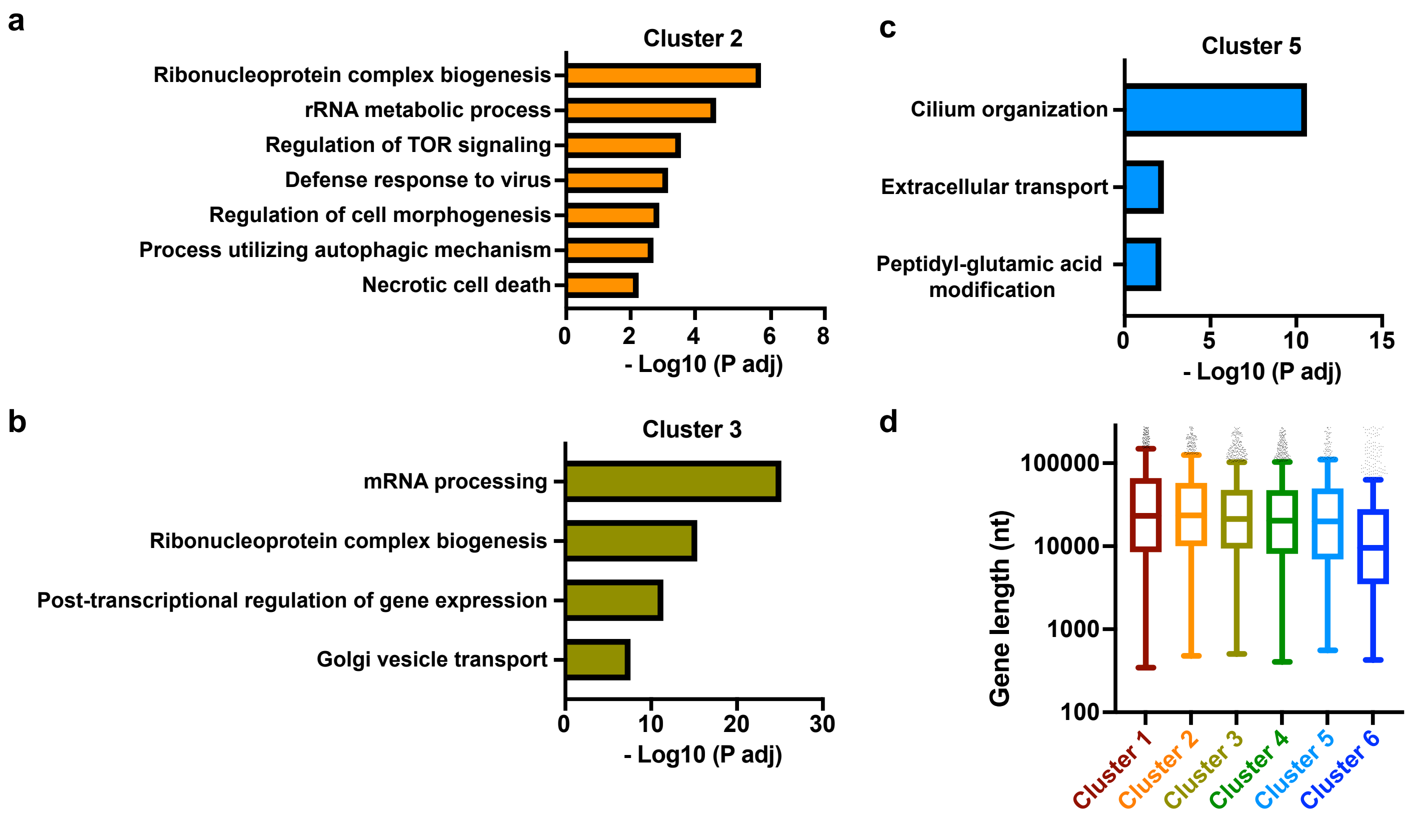


**Supplementary Fig. 1. Gene ontology terms and mRNA gene lengths for clusters of differentially expressed genes.**

**a-c.** Gene ontology terms and significance values for cluster 2 (**a**), cluster 3 (**b**), and cluster 5 (**c**) generated with ClusterProfiler and consolidated using Revigo.

**d.** Box plot representation of the distribution of gene length (TSS to TES) for mRNA genes in each cluster. Line represents median, box represents 25-75th percentile, whiskers represent 1.5X interquartile range.

Figure S2

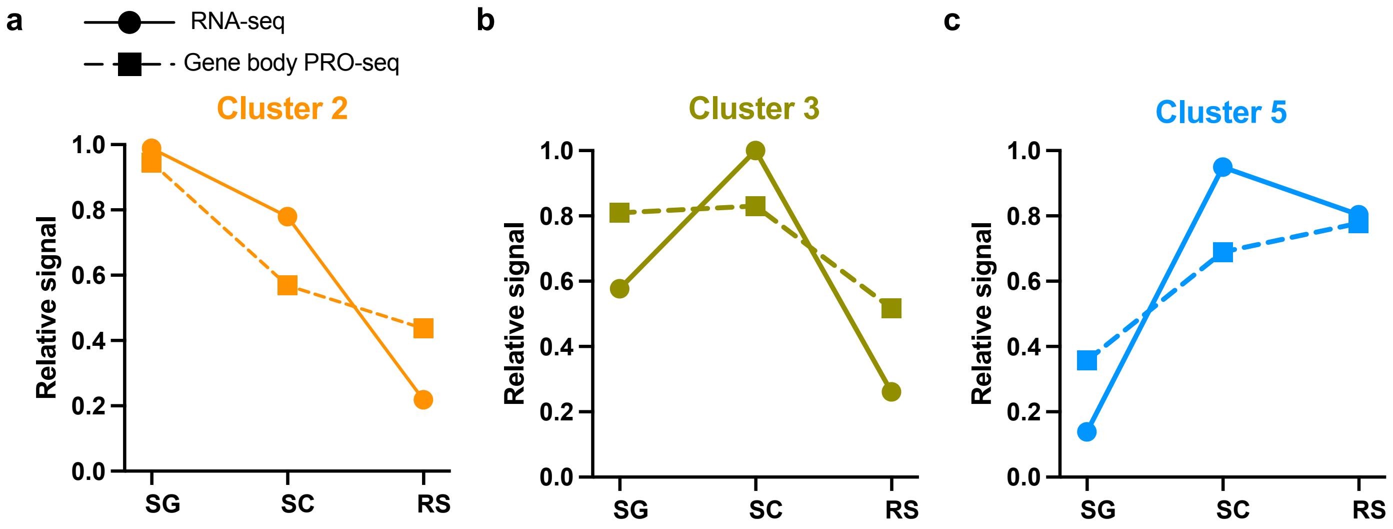


**Supplementary Fig. 2. Relative signal from RNA and PRO-seq gene body across cell types.**
**a-c.** Plots of average relative signal from RNA-seq (solid line and circle) or PRO-seq in gene bodies (dotted line and square) for each cell type, for clusters 2 (**a**), 3 (**b**), and 5 (**c**).

**Figure S3**

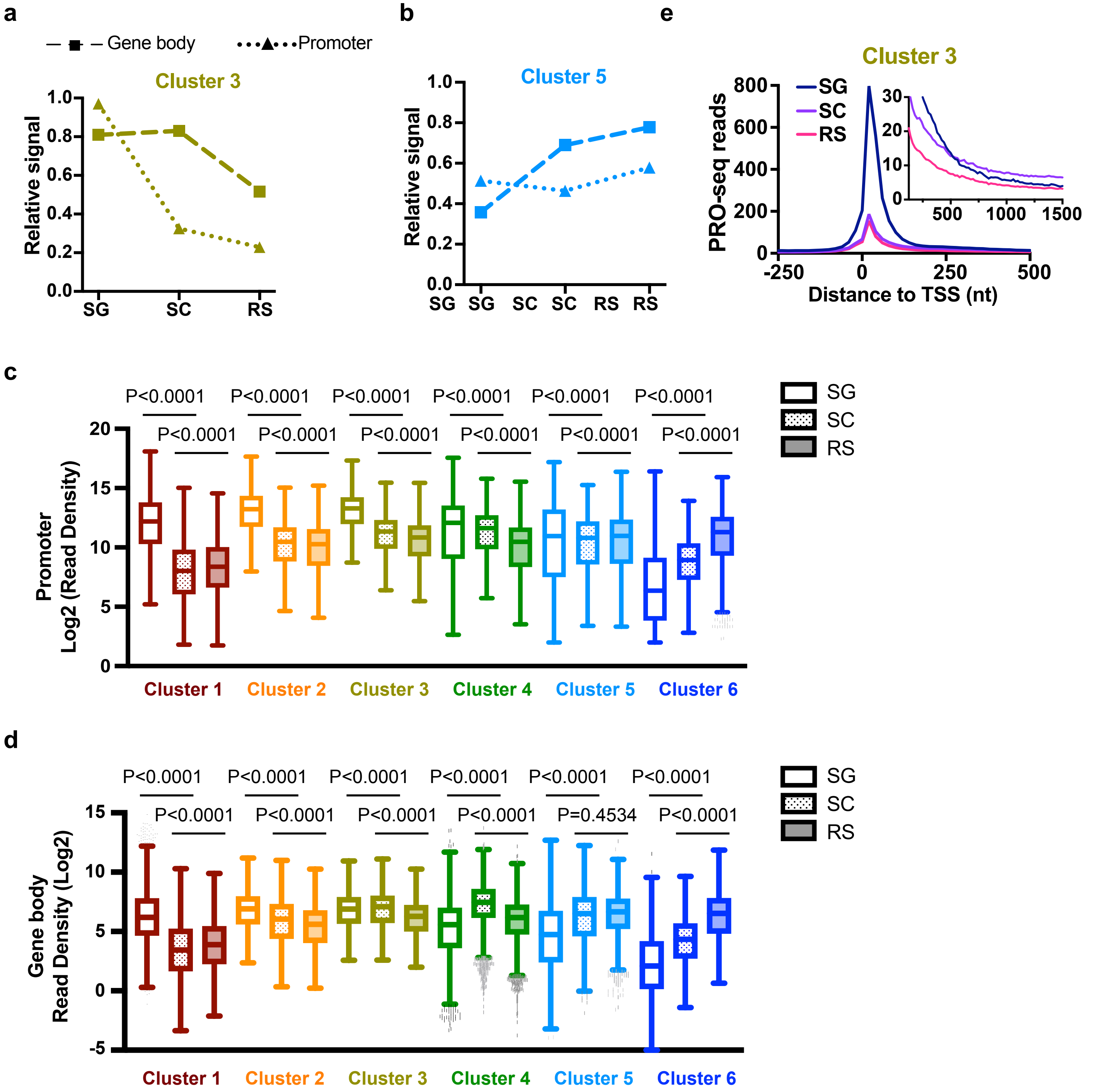


**Supplementary Fig. 3 Promoter and gene body counts by cluster and cell type.**

**a-b.** Plots of average relative signal from PRO-seq within promoter (dotted line with triangles) or gene body (dashed line with squares) regions for each cell type. Shown are data for genes in clusters 3 (**a**) 5 (**b**).

**c-d.** Box plot representations for the distribution of reads per kb in each cluster from PRO-seq promoter, (TSS to +150, **c**) or gene body (TSS+250 to TES, **d**), for each cell type. Legend indicates SG (empty), SC (dotted), and RS (shaded). P-values are from Wilcoxon matched-pairs test.

**e.** Metagene plot of average PRO-seq reads in each cell type for all protein-coding genes in cluster 3. Inset y-axis is zoomed-in to show gene body signal from +125 to +1500 nt downstream of the TSS.

### Figure S4

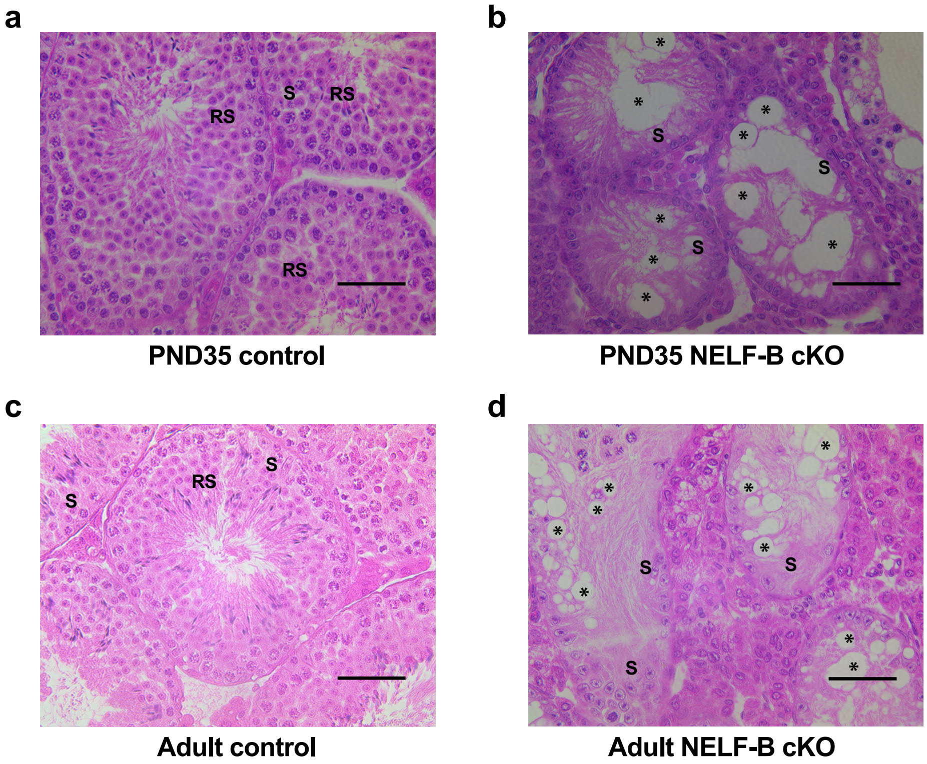


**Supplementary Fig. 4. NELF-B is required for the successful completion of spermatogenesis.**
**a-b.** H & E staining of testis cross sections of PND35 control (**a**) or NELF-B cKO (**b**) mice. NELF-B cKO testis show severe degeneration of the epithelium with vacuoles (asterisk) and only Sertoli cells (S) remaining in the epithelium compared to control mice which successfully generate round spermatids (RS). Scale bar, 50 µm.
**c-d**. H & E staining of testis cross sections of adult 9-month old control (**c**) or NELF-B cKO (**d**) mice. There was no recovery of spermatogenesis in the adult as shown by the vacuoles (asterisk) and lack of germ cells progressing to RS. Scale bar, 50 µm.

**Figure S5**

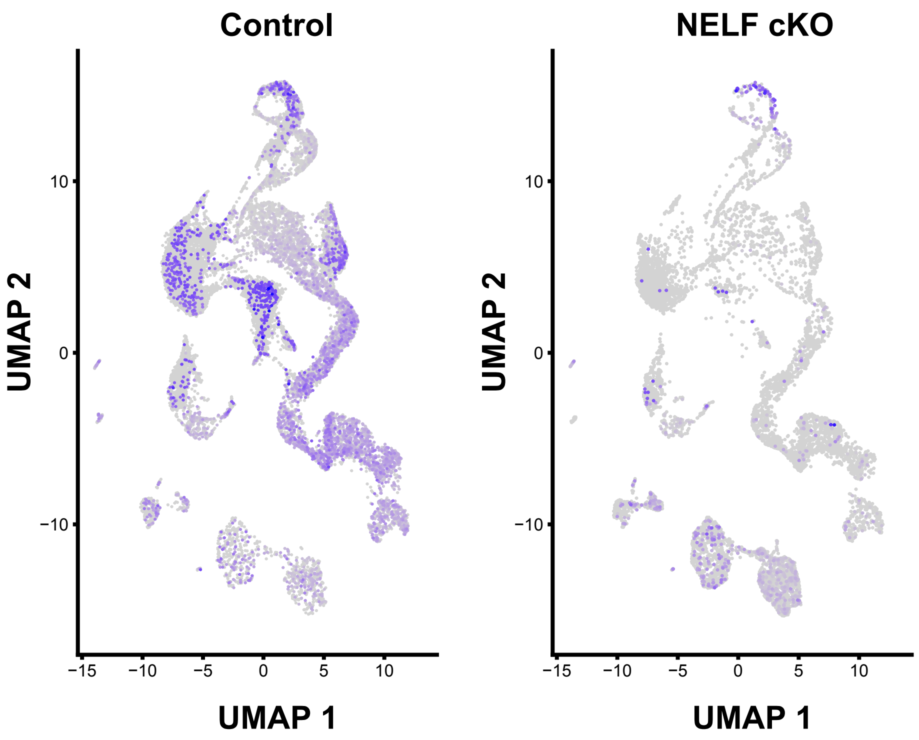


**Supplementary Fig. 5 NELF-B expression is selectively reduced in germ cells in NELF-B cKO mice, yet cells that reach RS stage often retain expression of NELF-B**

NELF-B expression across cell types in Control and NELF-B cKO mice, shown in purple. The bottom 1% of cells, in terms of the gene's expression, are not shaded.

**Figure S6**

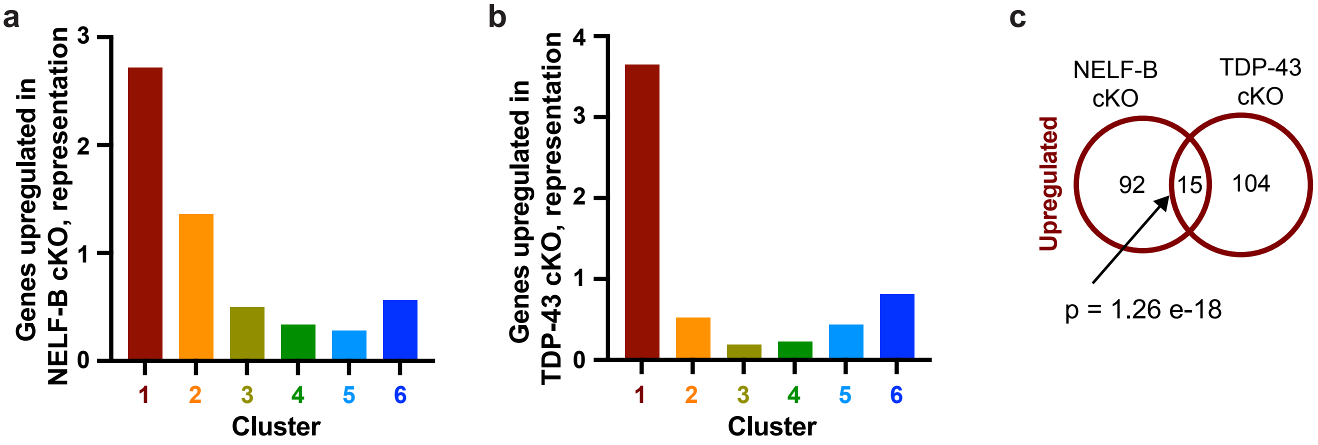


**Supplementary Fig. 6 Genes upregulated in SG in the absence of NELF-B or TDP-43 are over-represented in cluster 1 and have some overlap.**
**a-b.** Representation of upregulated genes from NELF-B cKO (**a**) or TDP-43 cKO mice (**b**), within each cluster (from Fig. 1f). A value of 1 indicates representation expected by chance.
**c.** Overlap between genes upregulated in NELF-B and TDP-43 mice. P-value for the overlap of gene lists was calculated using the hypergeometric distribution with the phyper function in R.

**Figure S7**

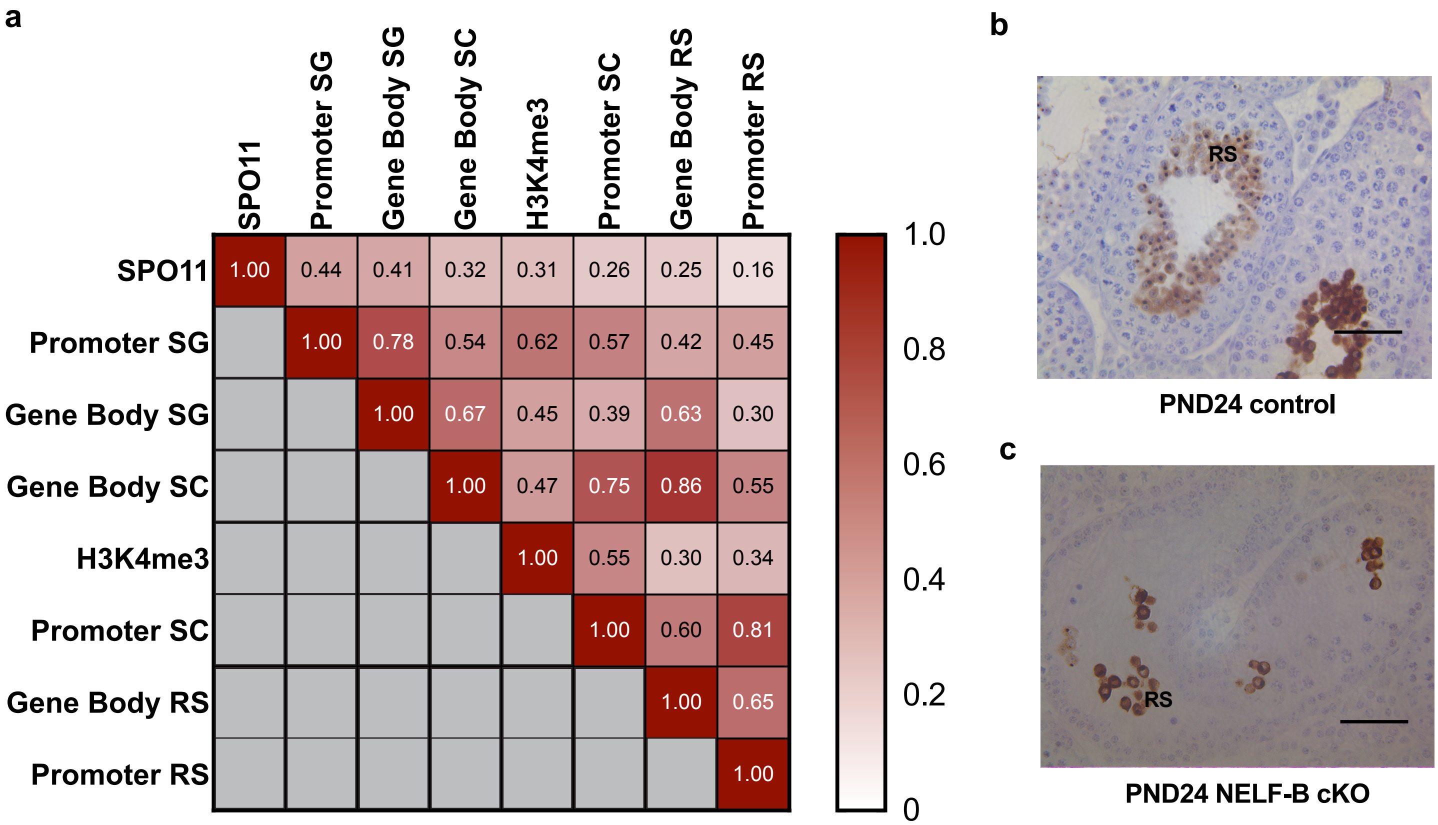


**Supplementary Fig. 7. SPO11 oligo reads indicative of DSBs correlate best with promoter PRO-seq signal in SG, and NELF-B cKO mice generate very few post-meiotic germ cells.**

**a.** Heatmap matrix of Spearman's correlation comparisons between SC SPO11 oligo sequencing (sum TSS+-500), H3K4me3 ChIP-seq (Sum TSS+-500), and PRO-seq promoter and gene body windows in SG, SC, and RS across all DE genes (n=17,078). Data sets are ordered left to right / top to bottom based on correlation with SPO11.

**b-c**. Immunohistochemistry of PND24 mouse testis using an antibody to the acrosomal marker SP-10. Control PND24 testis show the luminally arranged post-meiotic round spermatids (RS) with acrosome stained (**b**). PND24 NELF-B cKO testis show very few antibody-positive RS, indicating meiotic failure (**c**). Scale bar, 50 μm.
